# Hydroxychloroquine Proves Ineffective in Hamsters and Macaques Infected with SARS-CoV-2

**DOI:** 10.1101/2020.06.10.145144

**Authors:** Kyle Rosenke, Michael A. Jarvis, Friederike Feldmann, Benjamin Schwarz, Atsushi Okumura, Jamie Lovaglio, Greg Saturday, Patrick W. Hanley, Kimberly Meade-White, Brandi N. Williamson, Frederick Hansen, Lizette Perez-Perez, Shanna Leventhal, Tsing-Lee Tang-Huau, Martha Nason, Julie Callison, Elaine Haddock, Dana Scott, Graham Sewell, Catharine M. Bosio, David Hawman, Emmie de Wit, Heinz Feldmann

## Abstract

We remain largely without effective prophylactic/therapeutic interventions for COVID-19. Although many human clinical trials are ongoing, there remains a deficiency of supportive preclinical drug efficacy studies. Here we assessed the prophylactic/therapeutic efficacy of hydroxychloroquine (HCQ), a drug of interest for COVID-19 management, in two animal models. When used for prophylaxis or treatment neither the standard human malaria dose (6.5 mg/kg) nor a high dose (50 mg/kg) of HCQ had any beneficial effect on clinical disease or SARS-CoV-2 kinetics (replication/shedding) in the Syrian hamster disease model. Similarly, HCQ prophylaxis/treatment (6.5 mg/kg) did not significantly benefit clinical outcome nor reduce SARS-CoV-2 replication/shedding in the upper and lower respiratory tract in the rhesus macaque disease model. In conclusion, our preclinical animal studies do not support the use of HCQ in prophylaxis/treatment of COVID-19.

**One Sentence Summary:** Hydroxychloroquine prophylaxis/treatment showed no beneficial effect in SARS-CoV-2 hamster and macaque disease models.

---

Severe acute respiratory syndrome coronavirus 2 (SARS-CoV-2) is the causative agent of coronavirus disease 2019 (COVID-19) (*1*). SARS-CoV-2 infections were initially reported in China near the beginning of December 2019 (*2*). Following early spread through Asia, and subsequently to European, American and African countries, the virus is responsible for the third pandemic of the 21^st^ Century. With currently over 6.6 million confirmed cases and >390,000 deaths worldwide, health systems are stretched beyond limit with largely no proven treatment or prophylaxis available to reduce the burden (*3*). Public health measures combined with increasingly severe restrictions on public life have been implemented in many countries to stop SARS-CoV-2 transmission. The goal of current public health strategies is to flatten the epidemiologic SARS-CoV-2/COVID-19 curve to ease the burden on health care systems challenged by the highly intensive care required for a significant proportion of COVID-19 cases. Over 1,000 clinical trials are currently open or being established in different countries testing drugs such as lopinavir/ritonavir, dexamethasone, hydroxychloroquine (HCQ) and inhaled interferon beta-1a (*4*). Yet, many of these treatments have not been empirically tested in relevant SARS-CoV-2 animal disease models to determine preclinical efficacy, and thereby provide valuable insight into prioritization of drugs to move forward in humans.

At the time this work was started, the US FDA had given emergency approval for the use of chloroquine and HCQ in COVID-19 patients (*5*). *In vitro* data on the inhibitory effect of chloroquine and HCQ on SARS-CoV-2 replication had been published (*6-8*) and HCQ alone or in combination with the macrolide antibiotic azithromycin had been used in early clinical trials to treat COVID-19 cases with varying effect (*9-11*). Despite ongoing clinical trials, preclinical efficacy data on the effect of HCQ in SARS-CoV-2 animal disease models were lacking. Herein, we assessed the efficacy of HCQ prophylaxis and treatment in two established animal disease models, the Syrian hamster and rhesus macaque (*12, 13*).

First, we confirmed the *in vitro* inhibitory effect of HCQ on SARS-CoV-2 replication in Vero E6 cells. Cells were pretreated with differing drug concentrations and the effect on viral RNA load in tissue culture supernatant was determined 72 hours after infection by quantitative reverse transcriptase polymerase chain reaction (qRT-PCR) (fig. S1). The half-maximal effective concentration (EC_50_) value for HCQ was 164.7nM, consistent with low/sub-micromolar levels previously reported for the established *in vitro* inhibitory effect of HCQ on SARS-CoV-2 replication (*6-8*).

Having confirmed *in vitro* efficacy, we next tested the ability of HCQ to alter the course of SARS-CoV-2 in the Syrian hamster disease model (*12*). Five groups of hamsters (n=6 per group) were prophylactically or therapeutically treated with an intraperitoneal infection of a standard (6.5 mg/kg in PBS; human dose for malaria prophylaxis/treatment) or high (50 mg/kg in PBS) dose HCQ regimen; control groups were treated with vehicle only. Hamsters were intranasally infected with SARS-CoV-2 using a dose of 1×10^4^ median tissue culture infectious doses (TCID_50_). For prophylaxis, a single treatment was performed 24 hours prior to infection. The therapeutic treatment started 1 hour after SARS-CoV-2 infection and was continued for 3 consecutive days. Disease manifestation in this model is transient and clinical signs peak between days 3 and 5 post-infection with ruffled fur, increased respiration rate and reduced mobility (*12*). Virus replication and shedding was determined by qRT-PCR in swab samples (oral and rectal) collected on days 2 and 4, and lung tissue taken at necropsy on day 4 post-infection. Regardless of HCQ administration, all animals showed comparable high levels of genome copy numbers for oral swabs (>10^7^ genome copies/mL) and comparable lower numbers for rectal swabs (<10^6^ genome copied/mL) decreasing in all groups over time (Fig. 1, A and B). Like viral RNA loads in swabs, there was no significant difference in disease manifestation over the time of the study. Gross lung pathology was similar among the groups consisting of focally extensive areas of consolidation that failed to collapse upon removal (fig. S2). Viral lung loads on day 4 were high (10^14^ genome copies/g) but indistinguishable between all groups (Fig. 1C). Lung to body weight ratios were similar in all animals with no significant difference between groups (Fig. 1D). Overall, HCQ administered either as prophylaxis or treatment at standard or high doses did not have any significant impact on SARS-CoV-2 replication and shedding, nor disease manifestation and progression in the Syrian hamster model.

**Figure 1:**
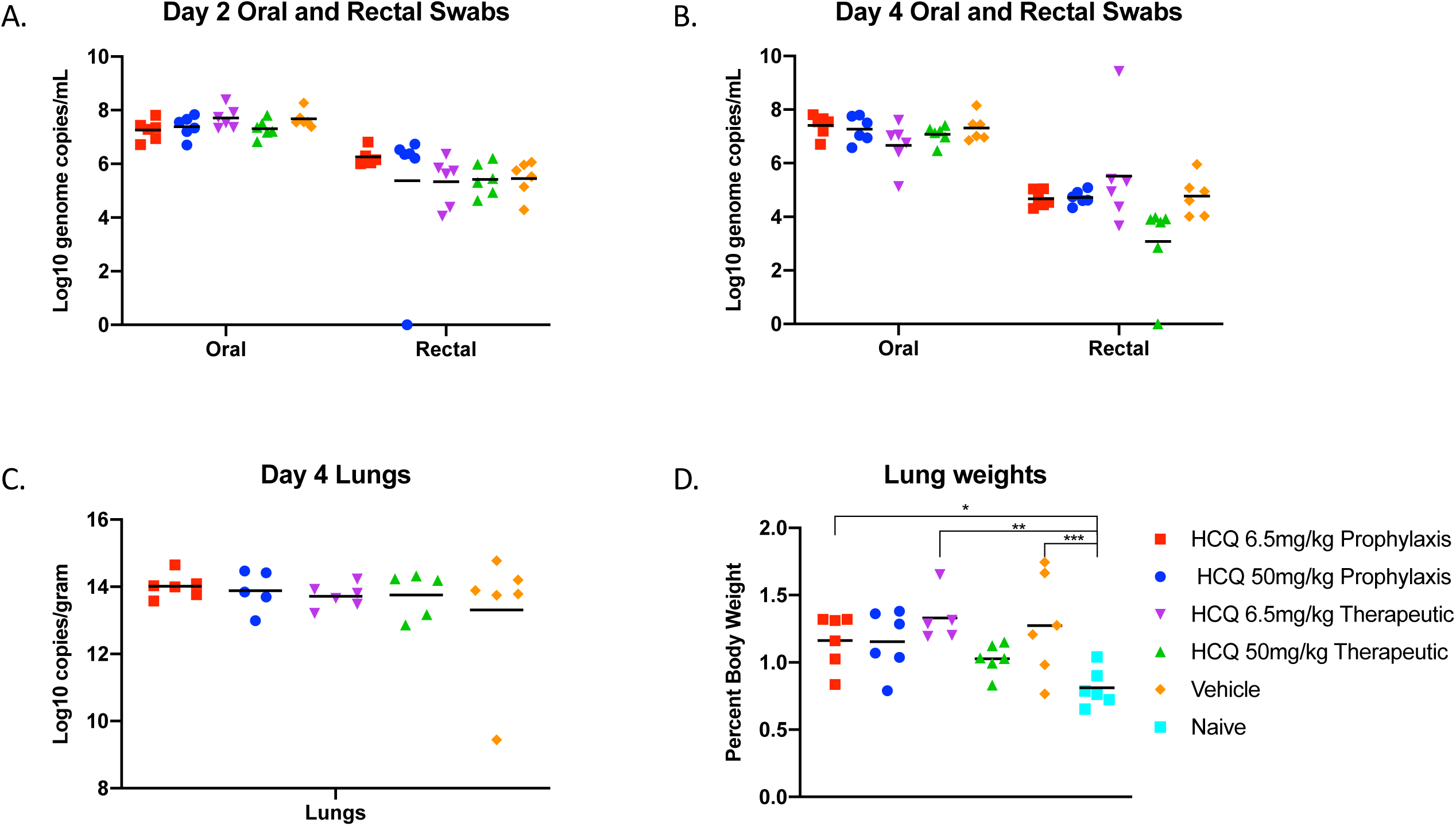
Syrian hamster model - viral shedding, viral load and pathology. Hamsters were infected with SARS-CoV-2 by the intranasal route. HCQ was administered either prophylactically one time at 24 hours prior to infection (6.5mg/kg and 50mg/kg) or treatment started 1 hour post-infection for 3 consecutive days (6.5mg/kg and 50mg/kg). Hamsters were scored for clinical signs daily and swabs (oral and rectal) were collected on day 2 and 4. Animals were euthanized on day 4 and lungs were harvested for pathology and virology. Swab and lung loads were determined by qRT-PCR. (A and B) Viral shedding. Oral and rectal swabs from day 2 and 4 were analyzed for viral genome copies by qRT-PCR. Swabs were analyzed as a correlate for viral shedding. (C) Viral load in lung tissue. Lung viral loads (genome copies) were determined by as a correlate for lower respiratory tract infection. No statistical significance was found among the groups presented in parts (A) to (C). (D) Lung to body weight ratio. Lung to body weight ratio was determined as an indicator for pneumonia with lung edema. Statistically significant differences were only found when compared to lung to body weight ratios of naïve hamsters. Multiple t tests were used to analyze differences among groups.

Next, we assessed HCQ efficacy in the rhesus macaque; a recently developed nonhuman primate model displaying mild to moderate COVID-like disease upon SARS-CoV-2 infection (*13*). Similar to the hamster study, we investigated the effect of HCQ when administered either prophylactically or as a treatment after infection. For the prophylactic arm, 10 healthy rhesus macaques were randomly divided into vehicle control and HCQ prophylaxis groups (n=5 per group). Animals were treated by oral gavage with either vehicle (PBS) or HCQ (6.5mg/kg in PBS) three times one week apart (day -9, day -2 and day 5) (Fig. 2A). To test the efficacy of HCQ as a treatment, a separate group of 10 healthy rhesus macaques were randomly divided into vehicle control and HCQ treatment groups (n=5 per group). Animals were treated by oral gavage with either vehicle (PBS) or HCQ (6.5mg/kg in PBS) starting 12 hours post-infection followed by treatment at 18, 36, 60, 84 and 108 hours post-infection (Fig. 2B). Animals in all groups were infected on day 0 with SARS-CoV-2 (total dose 2.8 ×10^6^ TCID_50_ by a combination of four routes (intratracheal, oral, intranasal and ocular) as previously described (*13, 14*). Animals were monitored at least twice daily using an established scoring sheet designed to assess clinical signs of disease (*13,15*). Multiple physical examinations were performed on different days pre- and post-inoculation including a clinical evaluation, radiographs, blood collection, and swabs (oral and nasal). Bronchoalveolar lavage (BAL) was performed on days 3, 5 and 7 (post-mortem) (Fig. 2, A and B). The endpoint for both studies was day 7 post-infection, at which time all animals were euthanized and necropsied.

**Figure 2:**
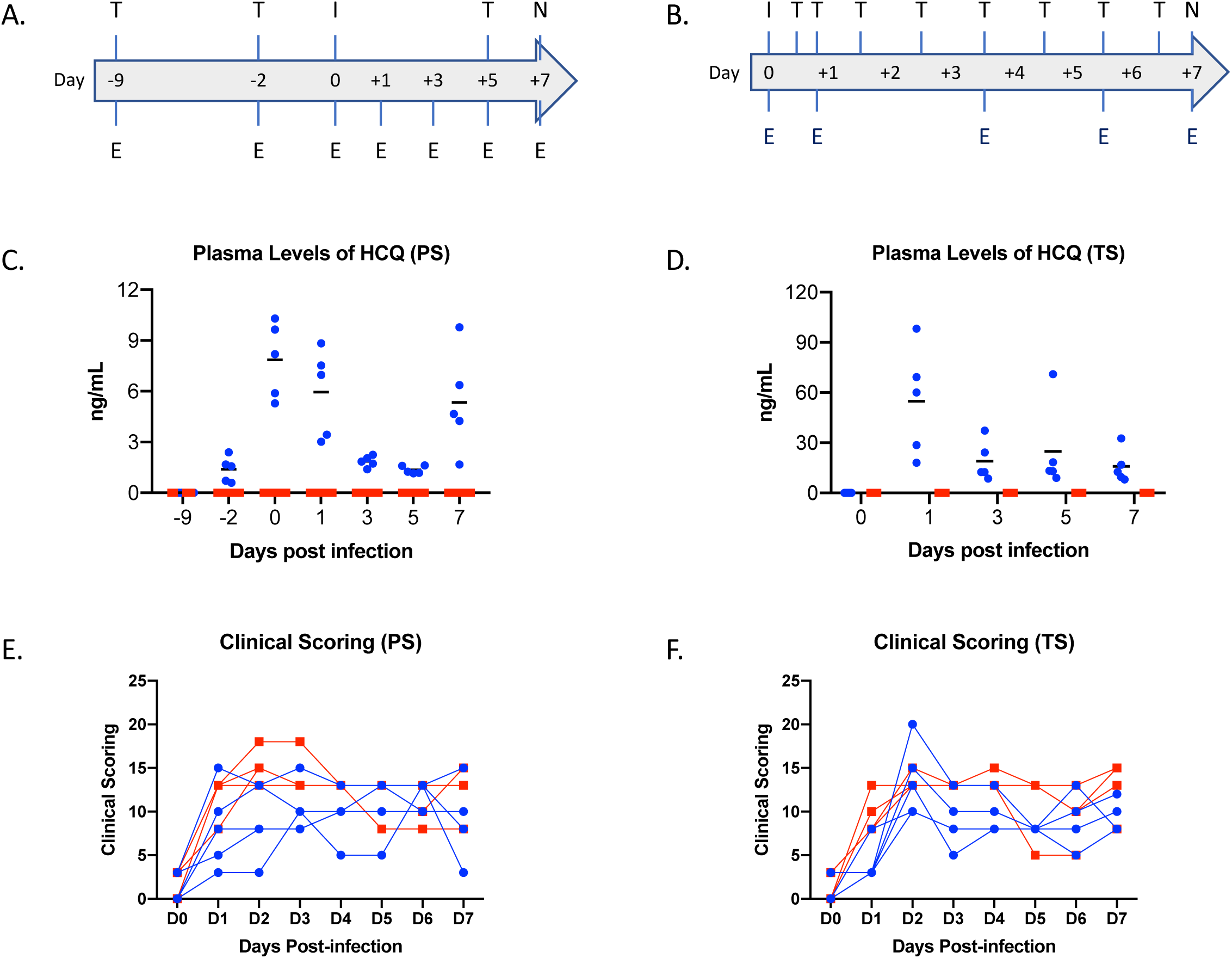
Rhesus macaque model – design, drug concentrations and clinical scoring. Macaques were infected with SARS-CoV-2 by the combined intratracheal, intranasal, oral and ocular routes. Animals were treated by oral gavage with either vehicle (PBS) or HCQ (6.5mg/kg in PBS). Administration was either one time per week for the prophylaxis arm or starting 12 hours post-infection followed by treatment at 18, 36, 60, 86 and 108 hours post-infection for the treatment arm. Animals were scored for clinical disease twice daily and examinations were performed as indicated. (A and B) Study design. The schematic depicts infection (‘I’), HCQ or vehicle treatment (‘T’) and examinations (‘E’). (C and D) Plasma levels of HCQ. HCQ levels were determined in both the prophylaxis and treatment study arms. Measurements reflect pre-dose levels of HCQ at each timepoint (limit of quantification = 0.5 ng/mL). (E and F) Clinical scores. Clinical scoring was performed twice daily by observation of non-anesthetized animals. The morning score is graphed here. Multiple t tests performed on individual days found no significance difference between groups. Area under the curve analysis was performed on each individual animal in each study. This analysis found a significant difference (p=0.004) between groups in the therapeutic study only. *Note:* red squares, vehicle-treated animals; blue circles, HCQ-treated animals; PS, prophylaxis; TS, treatment.

To ensure that drug was present in therapeutic quantities plasma levels of HCQ and its secondary metabolites were measured. HCQ was detected in plasma samples post-administration in all prophylactically or therapeutically treated animals with concentration ranging from 1.2 to 10.5ng/mL (3.6 nM to 31.3 nM) and 8 to 98 ng/mL (23.8 nM to 291.8 nM), respectively (Fig. 2, C and D). HCQ was also detected in lung tissue at time of necropsy in all prophylactically or therapeutically treated animals ranging from 0.85 to 4.18 ng/mg tissue and 1.39 to 11.54 ng/mg tissue, respectively. These numbers are in good agreement with the reported long half-life and large volume of distribution of HCQ (*16*). HCQ cytochrome p450 catalyzed secondary amine metabolites desethylchloroquine and desethylhydroxychloroquine, and the primary amine metabolite bisdesethylchloroquine are considered to be active forms of the drug in other disease models (*17*). Both desethylchloroquine and desethylhydroxychloroquine were detected in intermediate concentrations, while trace amounts of bisdesethylchloroquine were detected in both plasma and lung homogenate suggesting substantial persistence of active drug forms over the course of treatment (fig. S3). The plasma HCQ levels measured here fall within or near human therapeutically relevant ranges for other disease such as malaria and systemic lupus erythematosus (15 to 100 ng/mL plasma) (*18,19*). However, since SARS-CoV2 is a respiratory disease, levels of drug in lung tissue are a better indicator of therapeutic potential. Volume/concentration is difficult to estimate in tissue due to compartmentalization resulting in a non-homogenous distribution of the drug. However, using a water content of 80% by weight (*20*), day 7 levels in the lung indicated conservative estimates of at least 1 µg/mL (∼3.0 uM) in all animals, which is above the cell culture EC_50_ which we determined to be ∼ 0.2 uM (164.7 nM, 55 ng/mL here (fig. S1).

Macaques in both the prophylactic and treatment arms of the study first displayed clinical signs of SARS-CoV-2 infection on day 1, which peaked at day 2 and animals remained mildly to moderately ill until the study endpoint at day 7 (Fig. 2, E and F). Clinical signs included reduced appetite and ruffled fur followed by pale appearance and irregular increased abdominal respiration (table S1). Overall, animals in the vehicle treated groups appeared to have slightly higher clinical scores throughout, but daily differences were not statistically significant. Hematology and serum chemistry were unremarkable for all animals in both study arms. Radiographic signs in the prophylaxis, treatment and control groups were minimal over the study course (fig. S4). Pulmonary infiltrates, when seen, were noted to be of a mild unstructured interstitial pattern. The pattern was rarely seen in the upper lung, being more commonly found in middle and caudal lung lobes. No differences were noted in severity or appearance of radiographic signs between HCQ prophylaxis, treatment or control groups.

Nasal and oropharyngeal swabs were positive for SARS-CoV-2 RNA in all animals of both studies with the highest load on either day 1 or day 3, which then gradually decreased until the end of the study (Fig. 3, A – D). Viral load in nasal swabs were consistently higher than in oropharyngeal swabs. BAL samples were collected on days 3, 5 and 7 (post-mortem) and viral loads were similar to nasal and oropharyngeal swabs with decreasing loads over time (Fig. 3, E and F). Overall, there were no statistically significant differences in virus load and shedding between HCQ- and vehicle-administered animals in the prophylaxis and treatment regimens.

**Figure 3:**
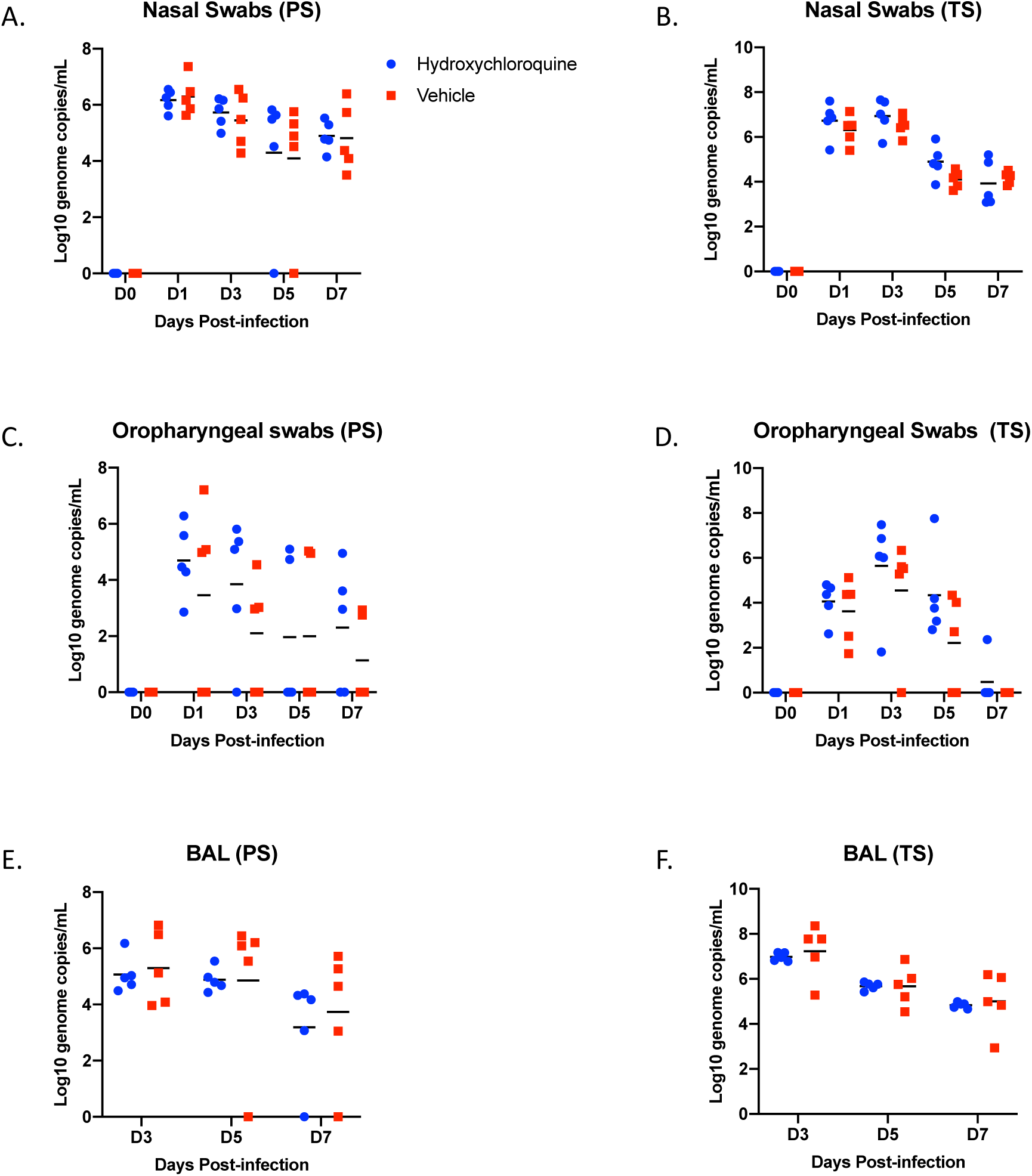
Rhesus macaque model – viral loads in lower and upper respiratory tract. Macaques were infected with SARS-CoV-2 as described in the legend of Figure 2. Swab samples (nasal and oropharyngeal) and bronchioalveolar lavage (BAL) were collected at all or indicated examination time points. Viral loads were determined by qRT-PCR as genome copies. (A and B) Nasal swabs. (C and D) Oropharyngeal swabs. (E and F) Bronchioalveolar lavage (BAL). No statistical significance was found among the groups presented in (A) to (F). Multiple t tests were used to analyze data and no significant difference was found. *Note:* red squares, vehicle-treated animals; blue circles, HCQ-treated animals; PS, prophylaxis; TS, treatment.

At necropsy, gross pathology revealed consolidated lungs in animals of all groups with lesions observed largely in the lower lung lobes, although some of the legions may have been the result of the post-mortem BAL (Fig. 4, A and B). All other gross pathology was normal except for enlarged cervical and mediastinal lymph nodes in several animals across the groups. Histological analysis of the lungs of animals in the different prophylaxis and treatment groups determined a comparable degree of pulmonary pathology when inoculated with SARS-CoV-2 similar to what had been published previously (*13,14*) (Fig. 4C). Lesions were mild to moderate and characterized as multifocal interstitial pneumonia frequently centered on terminal bronchioles. The pneumonia was evident by a thickening of alveolar septae by edema fluid and fibrin and small to moderate numbers of macrophages and fewer neutrophils. Infiltration of small numbers of pulmonary macrophages and neutrophils were noticed in alveoli. Lungs with moderate changes also had alveolar edema and fibrin with formation of hyaline membranes. There was minimal to moderate type II pneumocyte hyperplasia. Occasionally, bronchioles had necrosis, and loss and attenuation of the epithelium with infiltrates of neutrophils, macrophages and eosinophils. Perivascular infiltrates of small numbers of lymphocytes forming perivascular cuffs were noticed multifocally (Fig. 4C). Overall, there was no significant difference between vehicle and HCQ treated animals in either of the regimens, prophylaxis or treatment.

**Figure 4:**
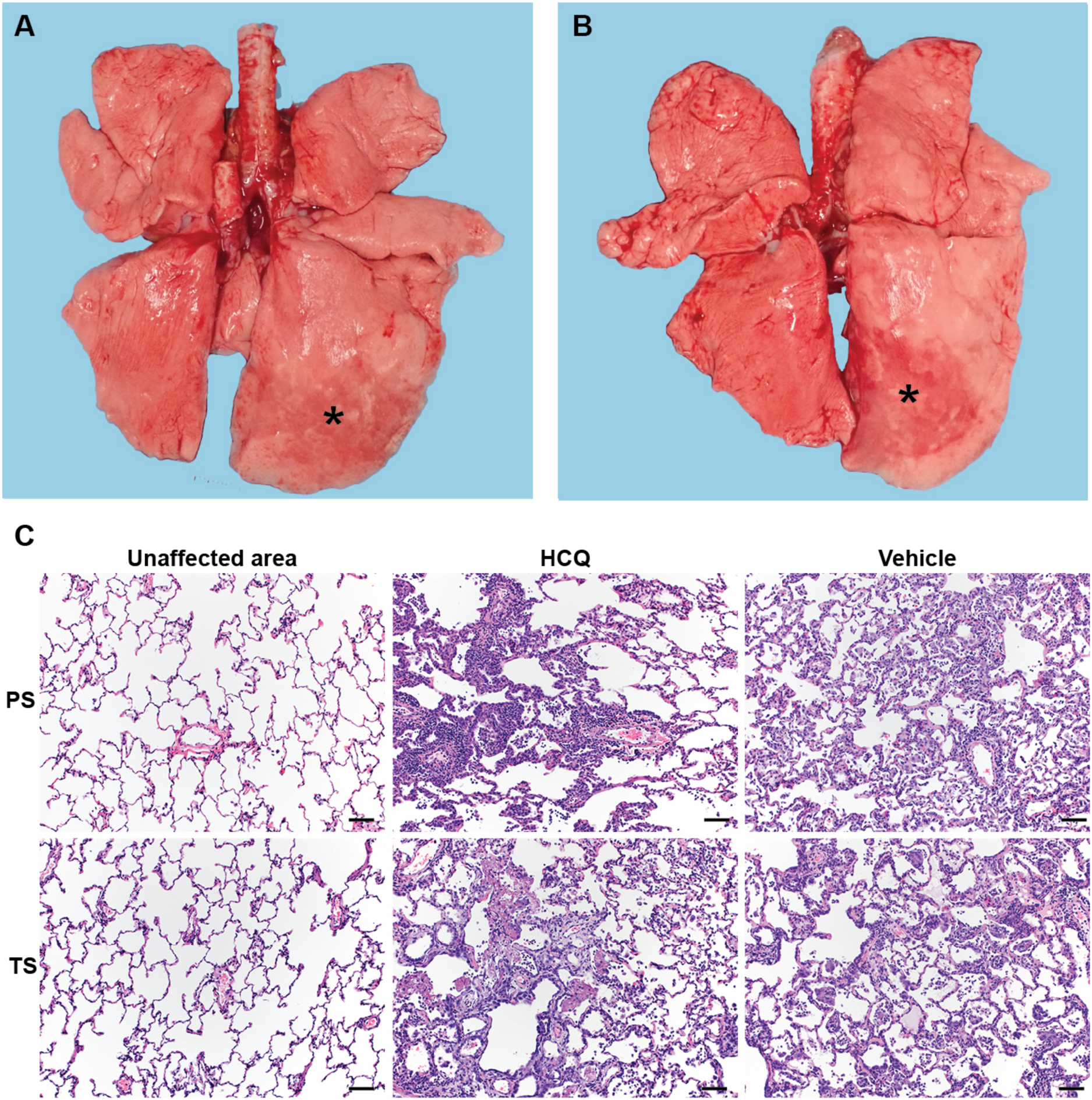
Rhesus macaque model – gross and histopathology. Macaques were infected with SARS-CoV-2 as described in the legend of Figure 2. Animals were euthanized on day 7 post-infection for gross pathology and histopathology. (A and B) Gross pathology with consolidated lower left lung lobe and area of post-mortem-BAL in the lower right lung lobe (asterisk). (C) Hematoxylin and eosin (H&E) staining revealed multifocal, minimal to moderate, interstitial pneumonia frequently centered on terminal bronchioles. Alveolar edema and fibrin with formation of hyaline membranes was only seen in lungs with moderate changes. Multifocal perivascular infiltrates of small numbers of lymphocytes that form perivascular cuffs. The left panels show areas of unaffected lung tissue. *Note:* PS, prophylaxis; TS, treatment.

Viral RNA loads were determined in several respiratory tissues using qRT-PCR (Fig. 5, A and C). Highest genome copy numbers were found in lung tissue with a marginal but not significant benefit for the HCQ-over the vehicle-treated group in the prophylaxis study arm when all lung lobe samples were combined (Fig. 5, B and D). Virus isolation from tissues was inconsistent among animals in the different groups, but at least one sample in each group showed infectious virus for almost all respiratory tissues (Fig. 5, A and C). There was no difference between animals of vehicle- and HCQ-treated groups in the prophylaxis and treatment study arms, which is consistent with the lack of any observed effect of HCQ on virus shedding parameters.

**Figure 5:**
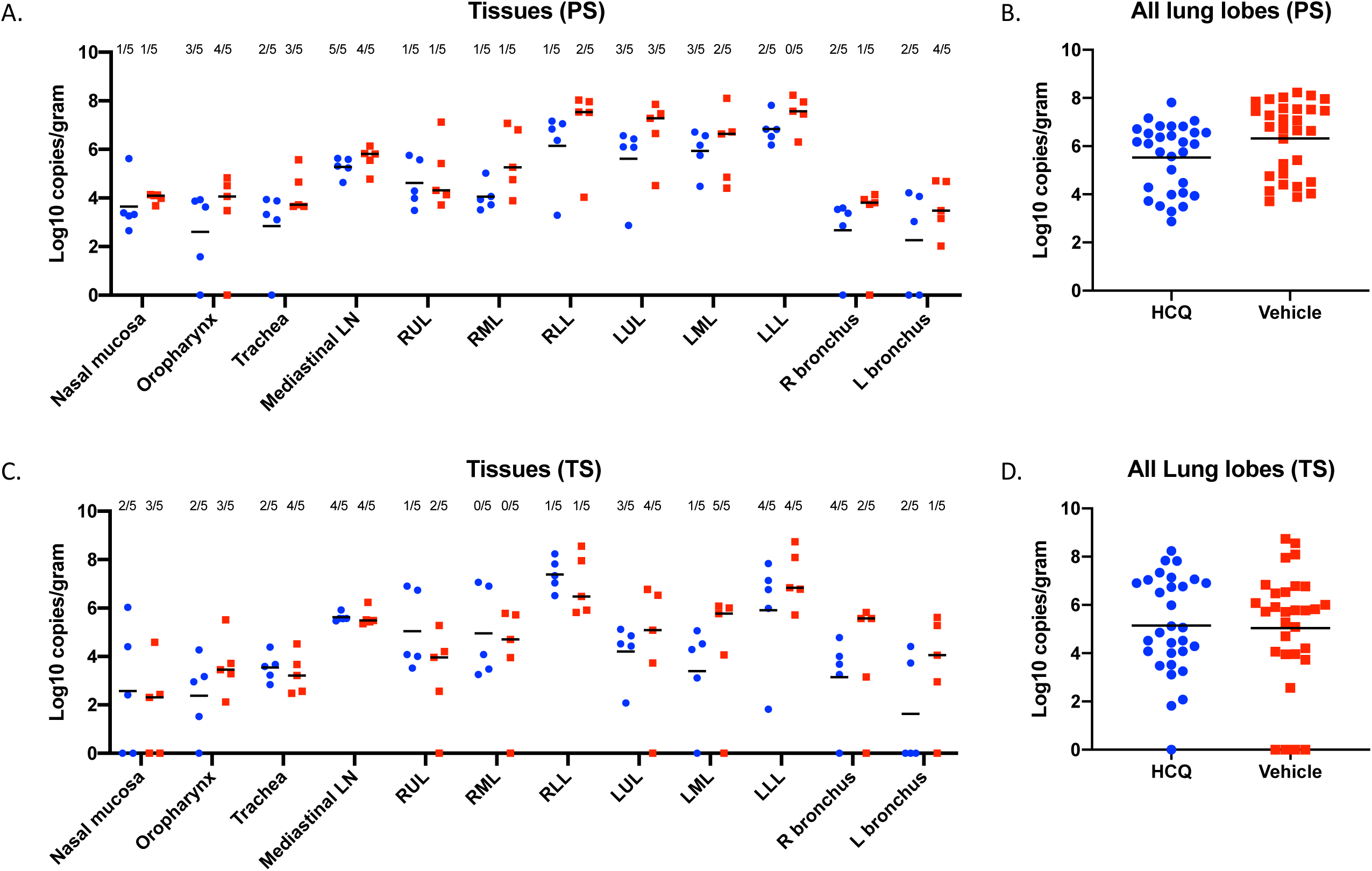
Rhesus macaque model – viral loads in respiratory tissues. Macaques were infected with SARS-CoV-2 as described in the legend of Figure 2. Animals were euthanized on day 7 post-infection for viral tissue load determination performed by qRT-PCR (genome copies) and virus isolation (infectious virus). (A) Viral loads in lower and upper respiratory tissues and mediastinal lymph nodes for the prophylaxis study arm (PS). Virus isolation is indicated in numbers on top (n/5). (B) Viral lung loads (PS). All lung lobe genome copy data were combined. (C) Viral loads in lower and upper respiratory tissues and mediastinal lymph nodes for the treatment study arm (TS). Virus isolation frequency (number of animals per group) is indicated at top (n/5). (D) Viral lung loads (TS). All lung lobe genome copy data were combined. No statistical significance was found among groups presented in parts (A) to (D). A linear model was used to analyze viral RNA levels in tissues and lung lobes. No significant difference was found between groups in either study. *Note:* red squares, vehicle-treated animals; blue circles, HCQ-treated animals; PS, prophylaxis; TS, treatment.

In this study we used two established COVID-like animal models (*12,13*) and applied the standard weight-based oral administration of HCQ prophylaxis and treatment of malaria in humans (*21*). For the Syrian hamster model, we also included a high HCQ dose regimen (7.5 times the standard dose regimen) both prophylactically and as a treatment. For prophylaxis we used a weekly dosing regimen. For treatment, we administered HCQ starting shortly after infection and continued daily until study end. HCQ pharmacokinetic studies in humans and animal models have demonstrated a rapid blood bioavailability following oral administration with peak levels being reached in 2 to 4 hours followed by rapid absorption in various tissues including the lung (*22,23*). Samples for drug pharmacokinetics in plasma were collected when the drug levels were low, just before the administration of the next treatment. Nevertheless, the measurements taken during both studies are in good agreement with data from humans and animal models and suggest accumulation of drug in the lung at therapeutic levels (*18,19*).

The use of HCQ and chloroquine as treatment options for COVID-19 patients may have been partially rooted in early observations for their effect in impairing SARS-CoV-2 replication *in vitro* (*6-8*). These *in vitro* studies, which we confirmed herein, identified HCQ (and other 4-aminoquinolines) as potent inhibitors of coronaviruses, including SARS-CoV-2, with low EC_50_ values within the range of antivirals such as remdesivir (*6*); a drug that is now approved for COVID-19 cases by the FDA. The mechanism of action of 4-aminoquinolones against SARS-CoV-2 *in vitro* is not well defined, but increasing endosomal pH, inhibition of autophagosome-lysosome fusion, impairment of enzymes important for virus replication, and effects on protein glycosylation have been proposed, which may result in interference with SARS-CoV-2 entry/fusion, replication and spread (*24, 25*). However, despite the promising *in vitro* effect observed by us and others, we did not observe any significant prophylactic or therapeutic benefit of HCQ following *in vivo* infection in two animal disease models.

The use of HCQ to treat COVID-19 has been controversial since the results of the first clinical trials (*9-11*). Nevertheless, HCQ has been promoted as a COVID-19 treatment option and became part of multiple recent large-scale clinical trials including one of four initial treatment options in the multinational WHO “Solidarity” clinical trial for COVID-19 (*26*). However, HCQ treatment does not come without risks as the 4-aminoquinolones are associated with multiple adverse effects such cutaneous adverse reactions, hepatic failure, and ventricular arrythmia; overdose is also difficult to treat (*21*). The US FDA recently updated its guidance by warning against use of HCQ outside of the hospital setting because of the potential for serious adverse effects (*27*). Over past weeks, several clinical trials, such as the WHO Solidarity study, have been stopped or have excluded HCQ arms due to a lack of evidence for therapeutic efficacy, and an increase level of adverse effects in COVID-19 patients (*26, 28, 29*). One influential study that had indicated a detrimental effect of HCQ in COVID-19 patients has subsequently been retracted by the authors due to their inability to confirm the veracity of the data (*29, 30)*, and the Solidarity HCQ arm has been resumed (*26*). Similarly, a multinational UK-based (COPCOV) HCQ prophylactic trial involving healthcare workers at high risk for SARS-CoV-2 infection was paused less than a week after starting due to safety concerns (*31*); the impact of the retraction on the status of this trial remains to be ascertained. Clearly, the effectiveness of HCQ to prevent or reduce infection and thereby impact the clinical course of COVID-19 remains highly contentious at this time.

In conclusion, HCQ prophylaxis and treatment had no beneficial effect in the two animal disease models tested. There is always the consideration as to what extent animal data can be extended to the situation in humans, but in general the nonhuman primate models are considered good indicators and the ultimate preclinical models before moving drugs into clinical trials. Independent of the safety issues associated with HCQ, the preclinical data presented here does not support HCQ and likely other 4-aminoquinolines as being either an effective prophylactic treatment to reduce SARS-CoV-2 infection or therapeutic for use in COVID-19 patients.

## Acknowledgements

This work was funded by the Intramural Research Program of the National Institutes of Allergy and Infectious Diseases (NIAID), National Institutes of Health (NIH), and partially funded through awards to The Vaccine Group Ltd, and the University of Plymouth. We thank Hillary Marston, Karyl Baron and Steven Holland (all NIAID, NIH) for helpful discussion and access to the drug. We are thankful to the animal caretakers and histopathology group of the Rocky Mountain Veterinary Branch (NIAID, NIH) for their support with animal related work, and Anita Mora (NIAID, NIH) for help with the display items.

## Disclaimer

The opinions, conclusions and recommendations in this report are those of the authors and do not necessarily represent the official positions of the National Institute of Allergy and Infectious Diseases (NIAID) at the National Institutes of Health (NIH). There were no conflict in interests identified for any individual involved in the study.

