## supplementary material for "Hydroxychloroquine Proves Ineffective in Hamsters and Macaques Infected with SARS-CoV-2"

File includes:

Materials and Methods

Supplementary Table s1

Supplementary Figure s1

Supplementary Figure s2

Supplementary Figure s3

Supplementary Figure s4

### Materials and Methods

**Biosafety and ethics.** Work with infectious SARS-CoV-2 was approved by the Institutional Biosafety Committee (IBC) and performed in high biocontainment at Rocky Mountain Laboratories (RML), NIAID, NIH. Sample removal from high biocontainment followed IBC-approved Standard Operating Protocols. Animal work was approved by the Institutional Animal Care and Use Committee and performed by certified staff in an Association for Assessment and Accreditation of Laboratory Animal Care International accredited facility. Work followed the institution's guidelines for animal use, the guidelines and basic principles in the NIH Guide for the Care and Use of Laboratory Animals, the Animal Welfare Act, United States Department of Agriculture and the United States Public Health Service Policy on Humane Care and Use of Laboratory Animals. Syrian hamsters were group housed in HEPA-filtered cage systems enriched with nesting material and were provided with commercial chow and water *ad libitum*. Nonhuman primates were single housed in adjacent primate cages allowing social interactions, in a climate-controlled room with a fixed light-dark cycle (12-hr light/12-hr dark). They were provided with commercial monkey chow, treats, and fruit twice daily with water *ad libitum*. Environmental enrichment consisted of a variety of human interaction, manipulanda, commercial toys, videos, and music. Hamsters and nonhuman primates were monitored at least twice daily.

**Virus and cells.** SARS-CoV-2 isolate nCoV-WA1-2020 (MN985325.1) was kindly provided by CDC and propagated once at RML in Vero E6 cells in DMEM (Sigma) supplemented with 2% fetal bovine serum (Gibco), 1 mM L-glutamine (Gibco), 50 U/ml penicillin and 50 μg/ml streptomycin (Gibco). The used virus stock was Vero passage 4, free of contaminations and confirmed to be identical to the initial deposited Genbank sequence (MN985325.1). Vero E6 cells were maintained in DMEM supplemented with 10% fetal calf serum, 1 mM L-glutamine, 50 U/mL penicillin and 50 μg/mL streptomycin.

**Syrian hamster study design.** The hamster study was designed with two arms, prophylaxis and therapeutic. Hamsters were divided into groups for either prophylaxis treatments or therapeutic treatments (n=6 per group). Two groups were treated one time with either a 6.5 mg/kg or 50 mg/kg 24 hours prior to infection for the prophylaxis arm. There were two therapeutic groups, one group received 6.5mg/kg while a second received 50mg/kg. Treatments began 1-hour post-infection and were performed every 24 hours on days 1, 2 and 3 post-infection. A final group consisted of vehicle control animals that received the same volume of PBS as the prophylactic and therapeutic groups. All groups were infected intranasally with 100ul of 1x10^5 median tissue culture infectious dose ($TCID_{50}$) of SARS-CoV-2 (50 uL/nare). Animals were euthanized and samples collected at day 4 post-infection. All procedures were performed on anesthetized animals. Swabs (oral, rectal) were collected at days 2 and 4, and lung tissues were collected at necropsy on day 4 post-infection for pathology and virology.

**Rhesus macaque study design.** The study also consisted of two arms, prophylaxis and treatment (Fig. 1A). Animals were anesthetized for all procedures. For the prophylaxis arm, 10 healthy rhesus macaques (all male; 4.9 – 5.6 kg in weight) were randomly divided into vehicle control (n=5) and HCQ prophylaxis (n=5) groups. Animals were treated with either vehicle (PBS) or HCQ (6.5mg/kg in PBS) three times one week apart (days -9, day -2 and day 5) by oral gavage. In the second arm, 10 healthy rhesus macaques (all male; 5.7 – 7.3 kg in weight) were randomly divided into vehicle control (n=5) and HCQ treatment (n=5) group. Animals were treated with either vehicle (PBS) or HCQ (6.5 mg/kg in PBS) starting 12 hours post-infection followed by treatment at 18, 36, 60, 84 and 108 hours post-infection by oral gavage. All animals were infected on day 0 with a total dose of 2.8 x10^6 $TCID_{50}$ of SARS-CoV-2 by a combination of four routes (intratracheal, oral, intranasal and ocular) as established and recently used in a drug efficacy study. Animals were monitored at least twice daily using an established scoring sheet by the same person, who was blinded to the group assignments, throughout the study (*1*). Physical examinations were performed on days -9, -2, 0, 1, 3, 5, and 7 and included a clinical evaluation, radiographs, venous blood draw, and swabs (oral, nasal and rectal). Bronchoalveolar lavage (BAL) was performed on day 3, 5 and 7 post infection. The study endpoint was day 7. Following euthanasia, necropsies were performed, and gross lung lesions were scored by a board-certified veterinary pathologist blinded to the group assignment.

**Liquid chromatography and mass spectrometry.** LCMS grade water, methanol, acetonitrile and formic acid were purchased through Fisher Scientific. All synthetic standards for molecular analysis were purchased from Santa Cruz Biotechnology Inc. Levels of HCQ and secondary metabolites were determined using methodology established previously with modifications (*2*). Plasma and cleared lung homogenates were gamma-irradiated (2 megarads) prior to removal from biocontainment according to IBC-approved protocol. Plasma samples were prepared for small molecule analysis by diluting a 25 µL aliquot with 100 µL of 0.1% formic acid and 1 mL of acetonitrile on ice. Clarified lung homogenate samples were prepared by adding 1 mL of acetonitrile to a 250 µL aliquot of lung homogenate. Samples were vortexed and incubated at -20 °C for 2 hrs.  Samples were centrifuged at 16,000 x g and 4 °C for 20 min. The clarified supernatants (1 mL) were recovered and taken to dryness in a Savant™ DNA120 SpeedVac™ concentrator (Thermo Fisher). Samples were resuspended in 100 µL of 50% methanol, 50% water (v/v) and centrifuged as before. The supernatant was taken to a sample vial for LCMS analysis. Samples were separated by reverse phase chromatography on a Sciex ExionLC™ AC system. Samples were injected onto a Waters Atlantis T3 column (100Å, 3 μm, 3 mm X 100 mm) and eluted using a binary gradient from 25 % methanol, 0.1 % formic acid to 100 % methanol formic acid over 4 min. Analytes were measured using a Sciex 5500 QTRAP® mass spectrometer in positive mode. Multiple reaction monitoring was performed using previously established signal pairs for each analyte and signal fidelity was confirmed by collecting triggered product ion spectra and comparing back to spectra of synthetically pure standards (*2*). All analytes were quantified against an 8-point calibration curve of the respective synthetic standard prepared in the target matrix and processed in the same manner as experimental samples. Limits of quantification in plasma for all metabolites was 0.5 ng/mL. Limit of quantification in lung homogenate for all metabolites was 6 ng/mL. Apparent and sample data were filtered prior to weight normalization.

**Hematology and serum chemistry**. Hematology was completed on a Procyte DX (IDEXX Laboratories, Westbrook, ME, USA) and the following parameters were evaluated: red blood cells (RBC), hemoglobin (Hb), hematocrit (HCT), mean corpuscular volume (MCV), mean corpuscular hemoglobin (MCH), mean corpuscular hemoglobin concentration (MCHC), red cell distribution weight (RDW), platelets, mean platelet volume (MPV), white blood cells (WBC), neutrophil count (abs and %), lymphocyte count (abs and %), monocyte count (abs and %), eosinophil count (abs and %), and basophil count (abs and %). Serum chemistries were completed on a Vetscan VS2 Chemistry Analyzer (Abaxis, Union City, CA) and the following parameters were evaluated: glucose, blood urea nitrogen (BUN), creatinine, calcium, albumin, total protein, alanine aminotransferase (ALT), aspartate aminotransferase (AST), alkaline phosphatase (ALP), total bilirubin, globulin, sodium, potassium, chloride, and total carbon dioxide.

**Thoracic radiographs.** Ventro-dorsal and right/left lateral radiographs were taken of nonhuman primates on clinical exam days and scored for the presence of pulmonary infiltrates by two clinical veterinarians according to a standard scoring system (0: normal; 1: mild interstitial pulmonary infiltrates; 2: moderate pulmonary infiltrates perhaps with partial cardiac border effacement and small areas of pulmonary consolidation; 3: serious interstitial infiltrates, alveolar patterns and air bronchograms). Individual lobes were scored. Scores from the lobes were then totaled and recorded per animal per day.

**Virus load.** RNA was extracted from swabs and BAL using the QIAamp Viral RNA kit (Qiagen) according to the manufacturer's instructions. Tissues were homogenized in RLT buffer and RNA was extracted using the RNeasy kit (Qiagen) according to the manufacturer's instructions. For detection of viral RNA, 5 µl RNA was used in a one-step real-time RT-PCR E assay (*3*) using the Rotor-Gene probe kit (Qiagen) according to instructions of the manufacturer. In each run, standard dilutions of RNA standards counted by droplet digital PCR were run in parallel, to calculate copy numbers in the samples.

**Virus titration.** Virus isolation was performed on lung tissues by homogenizing the tissue in 1 mL DMEM using a TissueLyser (Qiagen) and inoculating Vero E6 cells in a 24 well plate with 250 µL of cleared and a 1:10 dilution of the homogenate. One hour after inoculation of cells, the inoculum was removed and replaced with 500 µL DMEM (Sigma-Aldrich) supplemented with 2% fetal bovine serum, 1 mM L-glutamine, 50 U/mL penicillin and 50 µg/mL streptomycin. Six days after inoculation, cytopathogenic effect was scored and the $TCID_{50}$ was calculated.

**Histopathology and immunohistochemistry.** Histopathology and immunohistochemistry were performed on rhesus macaque tissues. After fixation for a minimum of 7 days in 10 % neutralbuffered formalin and embedding in paraffin, tissue sections were stained with hematoxylin and eosin (H&E). Stained slides were analyzed by a board-certified veterinary pathologist.

**Statistical analyses.** Statistical analysis was performed in Prism 8 (GraphPad).

**References.**

(1) D.L. Brining et al., Thoracic radiography as a refinement methodology for the study of H1N1 influenza in cynomologus macaques (Macaca fascicularis). *Comparative Medicine* **60**, 389-395 (2010).

(2) Y.S. Chhonker et al., Simultaneous quantitation of hydroxychloroquine and its metabolites in mouse blood and tissues using LC–ESI–MS/MS: An application for pharmacokinetic studies. *Journal of Chromatography B* **1072**, 320-327 (2018).

(3) V.M. Corman *et al.,* Detection of 2019 novel coronavirus (2019-nCoV) by realtime RT-PCR. *Eurosurveillance* **25**, 2000045 (2020).

**Table s1: Clinical observations**

| Prophylaxis | Day 1-3 | Day 4-7 |
|---|---|---|
| HCQ | Reduced appetite, ruffled fur, irregular abdominal respirations<br>Score: 8/8/8 | Reduced appetite, ruffled fur, irregular abdominal respirations<br>Score: 10/13/13/8 |
| HCQ | Reduced appetite, ruffled fur, slightly increased irregular abdominal respirations<br>Score: 10/13/10 | Reduced appetite, ruffled fur, slightly increased irregular abdominal respirations<br>Score: 10/10/10/10 |
| HCQ | Reduced appetite, ruffled fur, slightly irregular respirations<br>Score: 5/8/8 | Reduced appetite, ruffled fur, slightly increased irregular abdominal respirations<br>Score: 1013/10/10 |
| HCQ | Reduced appetite, ruffled fur, slightly increased irregular respirations<br>Score:3/3/10 | Reduced appetite, ruffled fur, slightly increased irregular respirations<br>Score: 5/5/13/3 |
| HCQ | Reduced appetite, ruffled fur, increased irregular respirations, abdominal & chest ↑ day 2&3, very pale appearance, dehydrated, diarrhea<br>Score: 15/13/15 | Reduced appetite, ruffled fur, increased irregular respirations, abdominal & chest ↑ day 2&3, very pale appearance, dehydrated<br>Score: 13/13/15 |
| Vehicle | Reduced appetite, ruffled fur, increased respirations↑↑ day 2&3<br>Score: 13/13/13 | Reduced appetite, ruffled fur, increased respirations↑↑ day 5&7, very agitated<br>Score: 13/13/13/13 |
| Vehicle | Reduced appetite, ruffled fur, increased respirations↑↑ on day 2&3,<br>Score:13/15/13 | Reduced appetite, ruffled fur, increased respirations<br>Score: 13/13/10/15 |
| Vehicle | Reduced appetite, ruffled fur, increased irregular abdominal respirations↑↑ on day 2& 3,<br>Score: 13/15/13 | Reduced appetite, ruffled fur, increased irregular abdominal respirations↑↑ on day 4&5,<br>Score: 13/13/13/13 |
| Vehicle | Reduced appetite, ruffled fur, increased irregular abdominal respirations↑↑ on day 2& 3, diarrhea<br>Score: 8/15/13 | Reduced appetite, ruffled fur, increased irregular abdominal respirations↑↑ on day 4-7,<br>Score: 13/13/13/13 |
| Vehicle | Hunched posture, ruffled fur, increased abdominal respirations, decreased appetite, clear nasal discharge, flushed appearance<br>Score: 13/18/18 | Increased irregular abdominal respirations, reduced appetite<br>Score: 13/8/8/8 |

| Treatment | Day 1-3 | Day 4-7 |
|---|---|---|
| HCQ | Pale appearance, slow, increased abdominal respirations, reduced appetite<br>Score: 3/15/10 | Pale appearance, slow, increased irregular abdominal respirations, reduced appetite<br>Score:10/8/8/10 |
| HCQ | Increased abdominal respirations, reduced appetite<br>Score: 3/10/8/ | Increased abdominal irregular respirations↑↑ on day 4-6, extending his arms, reduced appetite<br>Score:8/8/10/12 |
| HCQ | Pale appearance, slow, increased abdominal respirations↑↑ on day 3, reduced appetite<br>Score: 8/10/8 | Pale appearance, slow, increased abdominal respirations↑↑ on day 4-7, reduced appetite<br>Score: 8/8/5/8 |
| HCQ | Reduced appetite, ruffled fur, irregular abdominal respirations, excessive salivation on day 2, red nose on day 3<br>Score: 3/13/5 | Reduced appetite, ruffled fur, increased irregular abdominal respirations<br>Score: 8/8/13/8 |
| HCQ | Reduced appetite, ruffled fur, increased irregular abdominal respirations, vomit<br>Score: 3/20/13 | Reduced appetite, ruffled fur, increased irregular abdominal respirations<br>Score: 13/8/5/8 |
| Vehicle | Pale appearance, ruffled fur, increased abdominal irregular respirations↑↑ on day 3, reduced appetite<br>Score: 8/15/13 | Pale appearance, ruffled fur, increased abdominal irregular respirations↑↑ on day 4-7, reduced appetite<br>Score: 15/13/10/13 |
| Vehicle | Reduced appetite, ruffled fur, increased irregular abdominal respirations↑↑ on day 3<br>Score: 8/13/13 | Pale appearance, slow, increased abdominal respirations↑↑ on day 4-7, reduced appetite<br>Score: 13/13/13/15 |
| Vehicle | Reduced appetite, ruffled fur, increased irregular abdominal respirations↑↑ on day 3<br>Score: 13/13/13 | Reduced appetite, ruffled fur, pale appearance, slow, increased irregular abdominal respirations↑↑ on day 3<br>Score: 15/13/13/15 |
| Vehicle | Reduced appetite, ruffled fur, increased irregular abdominal respirations<br>Score: 10/13/13 | Reduced appetite, ruffled fur, irregular abdominal respirations<br>Score: 13/5/5/8 |
| Vehicle | Reduced appetite, ruffled fur, slow, increased irregular abdominal respirations<br>Score: 8/15/13 | Reduced appetite, ruffled fur, slow, increased irregular abdominal respirations<br>Score: 15/13/10/15 |

HCQ, hydroxychloroquine; PS, prophylaxis; TS, treatment; The light gray underlined rows indicate animals that were administered drug either as prophylaxis or treatment.

### Supplemental Fig s1.

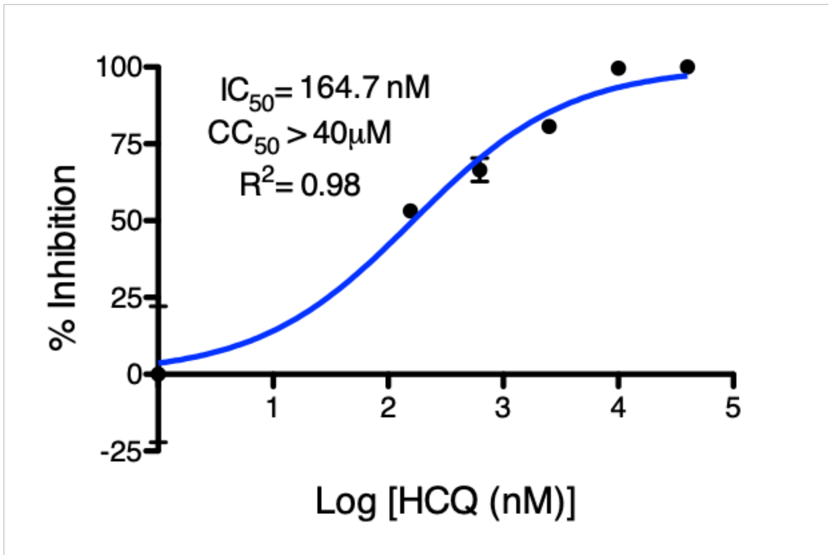

**Supplementary Figure s1: Half-maximal effective concentration ($EC_{50}$) of hydroxychloroquine (HCQ).** The $EC_{50}$ of HCQ was calculated based on inhibition of HCQ on SARS-CoV-2 replication at 72 hours in Vero E6 cells. Cells were pretreated for 1 hour with differing drug concentrations, followed by infection at a MOI of 0.01 for 1 hour. After 1 hour, media was replaced, and cells were cultured in the presence of drug for 72 hours at 37°C in a 5% $CO_2$ incubator. Virus yield in the cell supernatant was measured by quantitative RT-PCR of clarified culture supernatant. $EC_{50}$ values were determined following log-based transformation of drug concentrations and normalization to percentage inhibition based on diluent alone controls by fitting to drug-dose response curves using Prism software. Absence of toxicity at highest HCQ concentration (40uM) was confirmed using CellTiter-Glo® 2.0 Assay (Promega, Corp., Madison, WI, USA) as per manufacturer's protocol.

### Supplemental Fig s2.

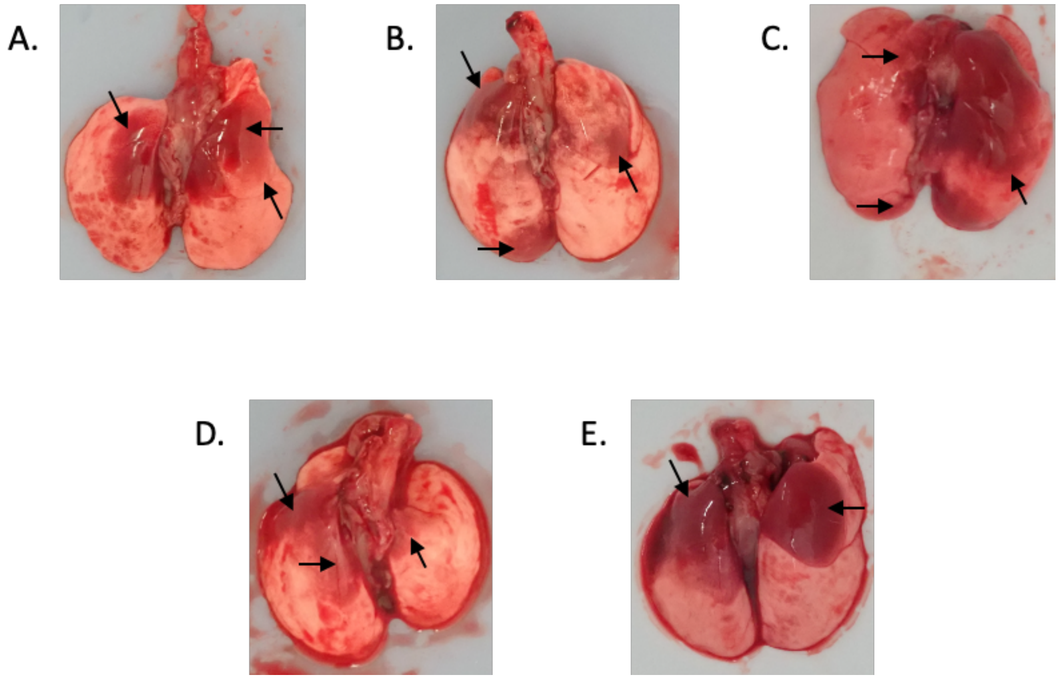

**Supplementary Figure s2: Syrian hamster model – gross pathology of lungs.** Syrian hamsters were infected with SARS-CoV-2 by the intranasal route. Hydroxychloroquine (HCQ) was administered either prophylactically (6.5mg/kg or 50mg/kg) or treatment (6.5mg/kg or 50mg/kg) or vehicle (equal volume) starting one hour post-infection followed by one daily treatment for 3 days. Animals were euthanized on day 4 post-infection for gross pathology and histopathology. Lungs were consolidated with large lesions observed largely in the upper lobes (representatively shown here one lung from one animal in each group. (A) 6.5mg/kg HCQ prophylaxis (PS). (B) 50mg/kg HCQ prophylaxis (PS). (C) 6.5mg/kg HCQ therapeutic (TS). (D) 50mg/kg HCQ therapeutic (TS). (E) vehicle groups. Note: PS, prophylaxis; TS, treatment.

Supplemental Fig s3.

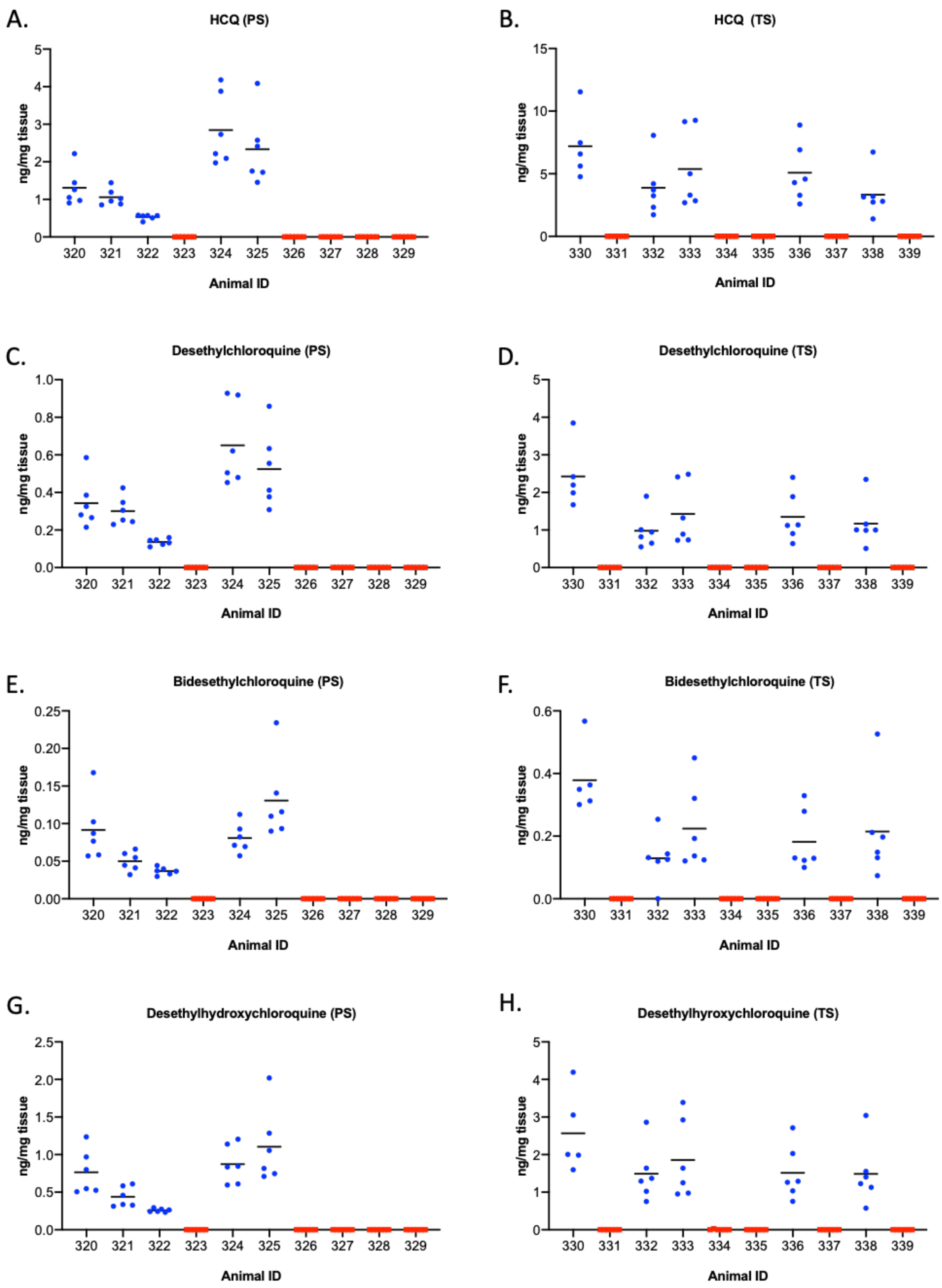

**Supplemental Figure s3: Levels of hydroxychloroquine (HCQ) and its metabolites desethylchloroquine, bisdesethylchloroquine and desethylhydroxychloroquine**.

Experimental dosing schedules in the lung tissue at day 7 post-inoculation for each animal were determined by LCMS. Each point represents a sample from a separate lobe of the lung. Mean ± SD are displayed. A single outlier was eliminated from display in animal 330 for all molecules measured using the ESD method and a significance level of 0.01. The limit of quantitation for all compounds was 0.5 ng/mL in the clarified homogenate prior to tissue weight normalization. **(A) HCQ (PS). (B) Desethylchloroquine (PS). (C) bisdesethylchloroquine (PS). (D) Desethylhydroxychloroquine (PS). (E) HCQ (TS). (F) Desethylchloroquine (TS). (G) bisdesethylchloroquine (TS). (H) Desethylhydroxychloroquine (TS).** Note: red square, vehicle treated; blue circle, HCQ, hydroxychloroquine; PS, prophylaxis; TS, treatment.

Supplemental Fig s4.

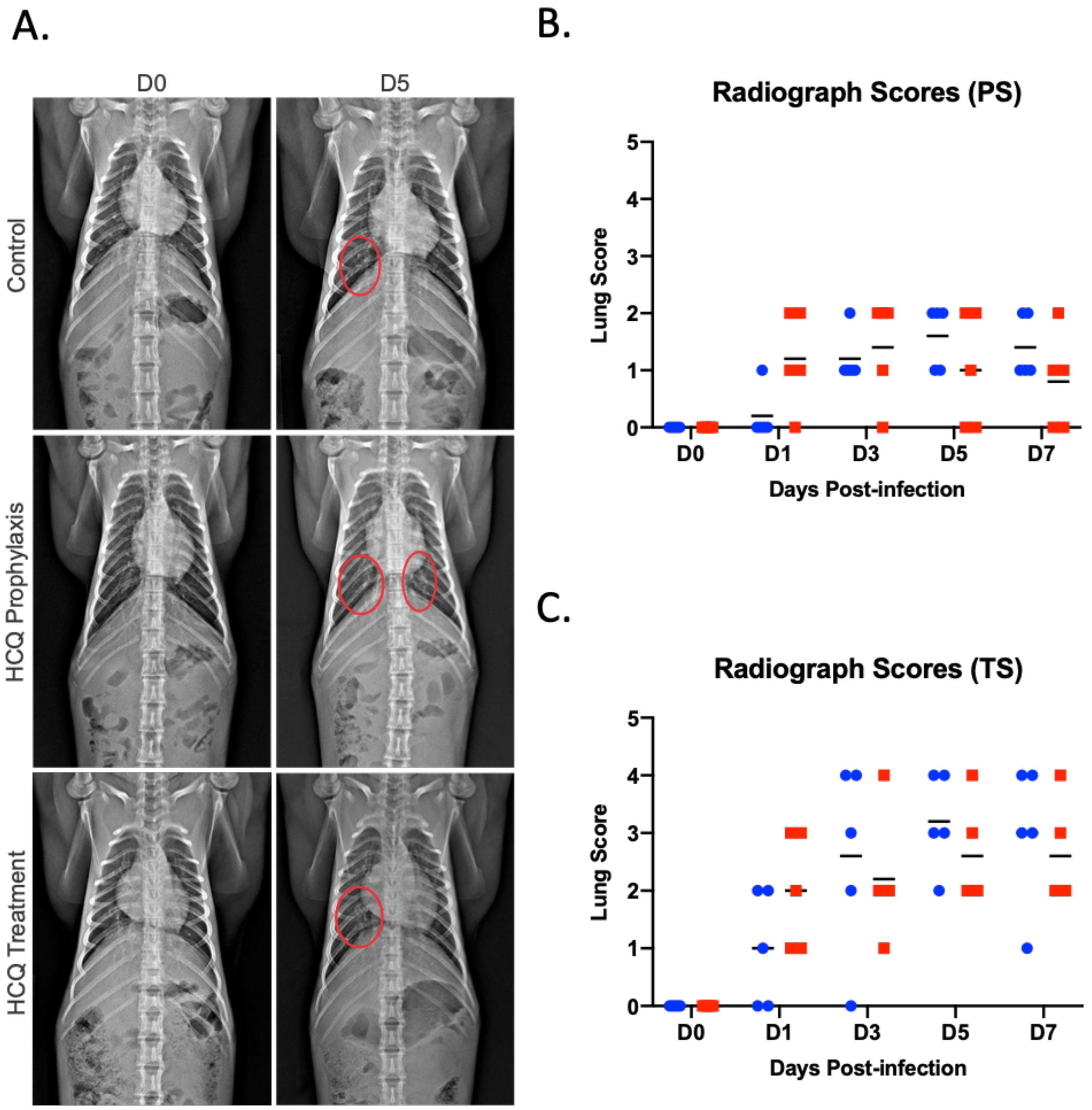

**Supplementary Figure s4: Rhesus macaque model – Radiography of lungs.** Ventro-dorsal and right/left lateral radiographs were taken of nonhuman primates on clinical exam days and scored for the presence of pulmonary infiltrates by two clinical veterinarians according to a standard scoring system (0: normal; 1: mild interstitial pulmonary infiltrates; 2: moderate pulmonary infiltrates perhaps with partial cardiac border effacement and small areas of pulmonary consolidation; 3: serious interstitial infiltrates, alveolar patterns and air bronchograms). Individual lobes were scored. Scores from the lobes were then totaled and recorded per animal per day. (A) Ventro-dorsal radiographs from representative animals in each group. Red ellipses indicate pulmonary infiltrates. (B) Radiograph scores – Prophylaxis (PS). (C) Radiograph scores – Therapeutic (TS). T tests were used to evaluate results, no significant difference was found between groups.
